# Evidence for circulation of Rift Valley fever virus in wildlife and domestic animals in a forest environment in Gabon, Central Africa

**DOI:** 10.1101/2023.10.30.564676

**Authors:** Pierre Becquart, Linda Bohou Kombila, Telstar Ndong Mebaley, Christophe Paupy, Déborah Garcia, Nicolas Nesi, Marie-Marie Olive, Jessica Vanhomwegen, Larson Boundenga, Illich Manfred Mombo, Camille Piro-Mégy, Matthieu Fritz, Léadisaelle Hosanna Lenguiya, Meriadeg Ar Gouilh, Eric M. Leroy, Nadine N’Dilimabaka, Catherine Cêtre-Sossah, Gael Darren Maganga

## Abstract

Rift Valley fever (RVF) is a mosquito-borne viral zoonosis caused by the RVF virus (RVFV) that can infect domestic and wild animals. Although the RVFV transmission cycle has been well documented across Africa in savanna ecosystems, little is known about its transmission in tropical rainforest settings, particularly in Central Africa. We therefore conducted a survey in northeastern Gabon to assess RVFV circulation among wild and domestic animals. Among 163 wildlife samples tested using RVFV-specific RT-qPCR, four ruminants belonging to subfamily Cephalophinae were detected positive. The phylogenetic analysis revealed that the four RVFV sequences clustered together with a virus isolated in Namibia within the well-structured Egyptian clade. A cross-sectional survey conducted on sheep, goats and dogs living in villages within the same area determined the IgG RVFV-specific antibody prevalence using cELISA. Out of the 306 small ruminants tested (214 goats, 92 sheep), an overall antibody prevalence of 15.4% (95% CI [11.5–19.9]) was observed with a higher rate in goats than in sheep (20.1% *versus* 3.3%). RVFV-specific antibodies were detected in a single dog out 26 tested. Neither age, sex of domestic animals nor season was found to be significant risk factors of RVFV occurrence. Our findings highlight sylvatic circulation of RVFV for the first time in Gabon. These results stress the need to develop adequate surveillance plan measures to better control the public health threat of RVFV.

**Author summary:** Rift Valley fever (RVF) is a mosquito-borne viral zoonosis caused by the RVF virus (RVFV) that can affect wild and domestic animals. Although the RVFV transmission cycle has been well documented across Africa in savanna ecosystems, little is known about its transmission in tropical rainforests, especially in Central Africa. We thus conducted a survey in northeastern Gabon to assess RVFV circulation among wild and domestic animals. In this study, we demonstrated for the first time in Gabon the presence of the RVFV in two wildlife species (Peter’s duiker *Cephalophus callipygus* and the blue duiker *Philantomba monticola*). In addition, we detected RVFV-specific antibodies in small domestic ruminants (sheep and goats) with an overall antibody prevalence of 15.4%, with a much higher seroprevalence rate in goats than sheep (20.1% versus 3.3%). Furthermore, RVFV-specific antibodies were also observed in a single (hunting) dog out of the 26 tested. These results stress the need to develop adequate surveillance plan measures to better control the public health threat of RVFV.

## Introduction

Among 175 human pathogenic species considered to be associated with emerging infectious diseases (EIDs), 75% are zoonotic — with many emerging over the past two decades in wildlife source [1]— making zoonotic EIDs a growing major threat to global health.

Although the emergence and re-emergence of diseases caused by arboviruses (viruses transmitted by arthropod vectors) is a constant concern in many African countries, their prevalence remains poorly documented due to the lack of efficient surveillance systems [2]. In addition, a significant number of vector-borne viruses are zoonotic, and there are gaps in the understanding of their ecology in natural wildlife niches and the factors that lead to their transmission to humans.

In Gabon, a total of 51 endemic or potentially endemic infectious viral diseases have been reported. Among them, 22 are of zoonotic origin and involve 12 families of viruses [3,4] with the most notorious arboviruses being Ebola, Marburg, and chikungunya, dengue, Rift Valley fever (RVF), yellow fever, West Nile fever and Zika. RVF is a World Organization for Animal Health (WOAH)-listed disease and a World Health Organization (WHO) priority disease for research and development due to its potential to cause major epidemics in humans [5]. RVF is a mosquito-borne, infectious disease caused by a negative single-stranded RNA virus named RVF virus (RVFV), a member of the *Phlebovirus* genus (family *Phenuiviridae*). In humans, RVFV infection is mostly pauci-symptomatic, but the illness can progress to hemorrhagic fever syndrome in few cases [6]. In animals, abortions and stillbirths in ruminants - domestic (cattle, sheep, goats and camels) or wild (buffaloes, antelopes, wildebeest) - resulting in major livestock deaths involving considerable economic losses in Africa, the Arabian Peninsula and the southwestern Indian Ocean region [7,8]. Epizootics (*i.e.* disease outbreaks that affect animals) of RVF are sporadic and often linked to persistent and heavy rainfall and flooding, which are in turn correlated with an abundance of mosquitoes of the *Aedes*, *Culex* and *Anopheles* genera, which are known to be involved in RVFV transmission [9,10]. Humans usually contract RVF through direct contact *via* aerosols of body fluid secretions of infected livestock and, to a lesser extent, may develop the disease through mosquito bites of infected mosquitoes [11].

Following the first description of RVFV in 1930 in Kenya [12], epizootics were recorded in East and South Africa until 1977; evidence from serological surveys (Angola, 1960; Cameroon, 1968; Chad, 1969) and virus isolations (Democratic Republic of Congo (DRC), 1936–1954; Central African Republic (CAR), 1969) have revealed contemporaneous circulation of RVFV in central Africa. Thereafter, the disease began to spread north to Sudan and Egypt, leading to the first massive epizootic/epidemic in Egypt in 1977–78, which affected 200,000 people and led to at least 600 deaths [13]. The disease was later recorded in Madagascar in 1979 and then West Africa (Senegal and Mauritania) in 1987 [14]. The epidemic potential and human health impact of this disease have been acutely felt on the African continent. RVF is enzootic/endemic in East and South Africa causing epizootics/epidemics in Egypt (2003), Kenya (2018), Somalia, Sudan, Madagascar (2008–2009, 2019–2021), South Africa (2009– 2011), Uganda (2016, 2023) [15] and various parts of West Africa, with inter-epizootic RVFV circulation. In 2000–2001, the virus left the African continent for the first time, reaching the Arabian Peninsula (Saudi Arabia, Yemen).

In Central Africa, at the crossroads of major African geographical regions experiencing RVF epizootics/epidemics, several studies have demonstrated the circulation of the virus in domestic ungulates as well as in humans in a savanna-type ecosystem in Cameroon, Gabon, Equatorial Guinea and the DRC [16–25], but no major epidemics or epizootics have been reported there, in contrast to East and South Africa, West Africa and Egypt. Nevertheless, little is known about RVFV in the tropical forests of Central Africa, with only a few serological surveys suggesting RVFV circulation. These surveys revealed the presence of RVF antibodies in antelopes, wild buffaloes, warthogs and elephants in CAR [26] and in the rural human population in Gabon [27]. Moreover, in southern Cameroon, the sylvatic circulation of RVFV was suggested to explain the presence of antibodies in locally bred goats [28]. Nonetheless, he sylvatic cycle of RVFV remains poorly documented in Central African rainforests. Several wildlife vertebrate hosts, particularly wild ungulates, are possibly involved in RVFV circulation involving forest mosquito species (belonging to genera *Aedes*, *Anopheles* and *Culex*) that are involved in, or are closely related to, domestic cycles. To date, little is known about RVFV sylvatic vectors in the forests of Central Africa, and the virus has only been isolated once in *Aedes* mosquitoes belonging to the *Neomelaniconion* subgenus and the *palpalis* species group collected in the CAR [29]. Moreover, isolation of RVFV from humans [30] together with serological RVFV evidence from Pygmy populations [31] suggest the existence of an RVFV forest cycle in the CAR and probably throughout Central Africa.

The study conducted here in Gabon was therefore intended to extend our knowledge of the sylvatic circulation of the RVFV in rainforests of Central Africa by investigating wildlife and domestic animals at the edge of rainforest. We demonstrated for the first time in Gabon the presence of the RVFV in two wildlife species (Peter’s duiker *Cephalophus callipygus* and blue duiker *Philantomba monticola*), along with RVFV-specific antibodies in livestock small ruminants and dogs.

## Materials and Methods

### Study area

The study was carried out in 19 villages located in the Zadié Department, located in the Ogooué-Ivindo province, northeastern Gabon. This area is mainly composed of primary tropical rainforests along three main routes radiating from Mekambo, the main city in Zadié: Mekambo-Mazingo (Route #1), Mekambo-Ekata (Route #2) and Mekambo-Malouma (Route #3) (Fig 1).

**Fig 1.**
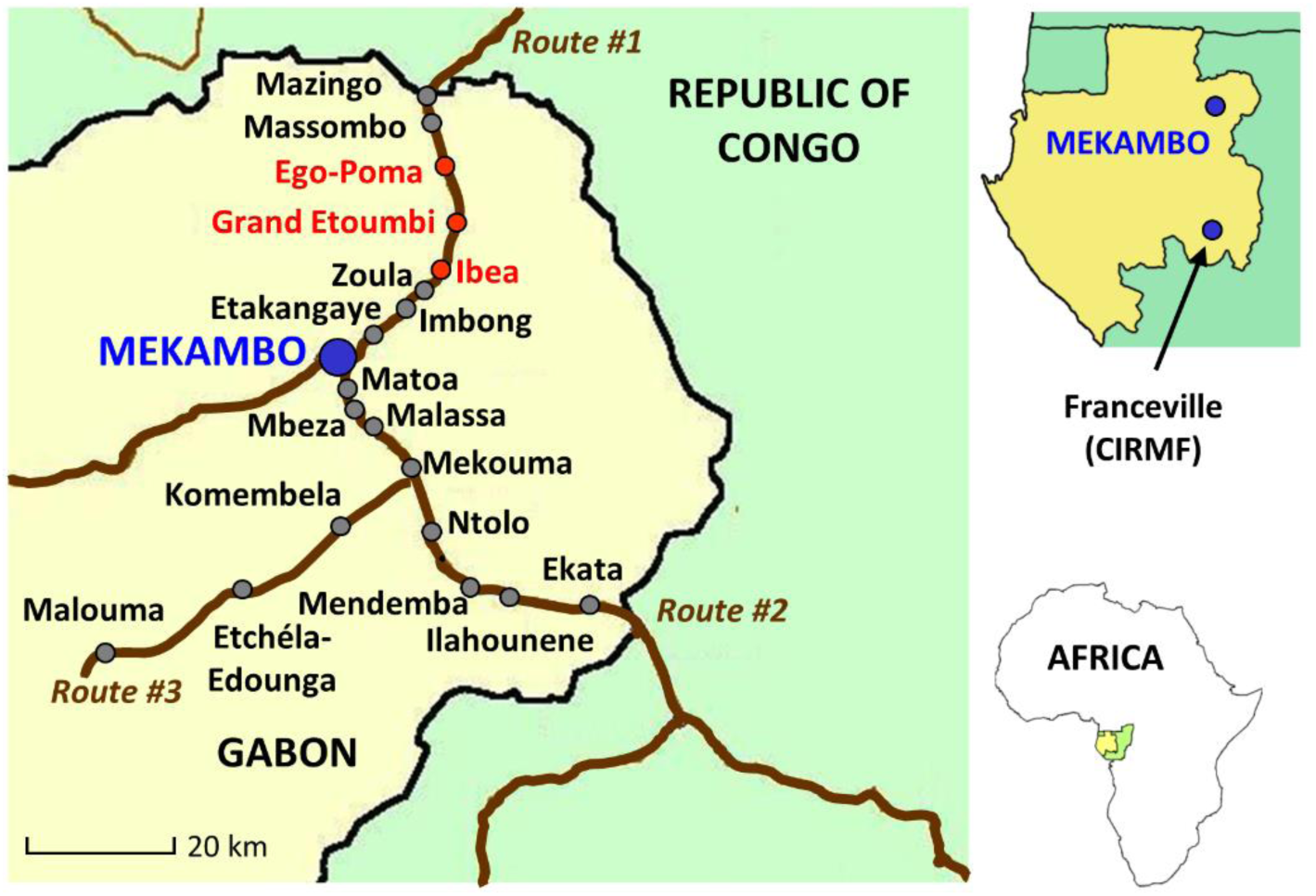
Map of the study area, Zadié Department, Gabon. Gray dots indicate the sampled villages and red dots, the villages where hunters brought back *Cephalophinae* detected positive for RVFV.

### Sampling and data collection

Wild animals were sampled along the three routes described above (Fig 1) in July 2019 during the dry season and legal hunting season. Organ samples (liver and spleen) were collected from animals hunted in the surrounding forest and displayed roadside and sold for consumption (Table 1). Samples were temporarily stored in liquid nitrogen at the Mekambo health center, before being transferred to CIRMF (*Centre Interdisciplinaire de Recherches Médicales de Franceville*) laboratory for storage at −80°C.

**Table 1.** List of wild animal species screened for the RVFV genome.

| **Wild animal species** | ***n*** |
| --- | --- |
| *Atherurus* sp. | 25 |
| *Cephalophus agilbigi* | 1 |
| *Cephalophus callipygus* | 4 |
| *Cephalophus dorsalis* | 13 |
| *Philantomba monticola* | 92 |
| *Cephalophus* sp. | 22 |
| *Cercopithecus cephus* | 1 |
| *Genetta abyssinica* | 2 |
| *Genetta* sp. | 2 |
| *Potamochoerus porcus* | 1 |
| **Total** | **163** |

Additionally, livestock (goats and sheep) and dogs were sampled at two time points, once in November 2018 (short rainy season) and once in July 2019 (long dry season), along Route #1 and Route #2 (Fig 1). There is no census of livestock in this region. Domestic animals were selected based on the willingness of the livestock owners to cooperate with the study. Thus, the number of sheep and goats sampled depended on the livestock owner’s availability and their ability to restrain their animals for sampling. Data on species, sex, period of sampling and age (or sexual maturity stage) were collected using a standard questionnaire submitted to each animal owner. Sheep and goats were classified as young or adult according to the criterion of sexual maturity: young (under 3 years old) and adult (aged >3 years) using both morphological characters observed by the veterinarians and information provided by animal owners. For each domestic animal, a blood sample was collected in EDTA tubes upon jugular venipuncture and preserved in a cooler box until transport to the laboratory.

### RVFV genome detection in wildlife

A total of 163 wild animals (comprising mostly *Cephalophus* spp. ruminants and *Atherurus* spp. rodents) were sampled (Table 1) and tested for the presence of RVFV genome using a RT-qPCR method. Briefly, after grinding up the organs (liver and spleen) in RA1 lysis buffer supplemented with a 1% Triton X-100 solution (Sigma, France), RNA was extracted using the Nucleospin RNA kit (Macherey-Nagel, Germany) followed by a RVFV-specific RT-qPCR amplification [32] in a Lightcycler L96 (Roche) equipment. When RVFV was detected in wildlife samples, DNA was extracted using the Qiagen DNeasy Blood & Tissue Kit (Qiagen, Courtaboeuf, France) in order to to amplify a 710 bp long fragment of the mitochondrial cytochrome oxidase I (*COX1*) gene using PCR to identify/confirm the vertebrate species [33]. *COX1* sequences generated were then aligned and compared with Cephalophinae sequences from central Africa.

### Sequencing and phylogenetic analysis of RVFV

All positive samples after the full-length RVFV S segment PCR amplification following the protocol defined in [34] were sequenced. The phylogenetic analyses were done after multiple alignments of the obtained sequences, along with GenBank reference sequences using ClustalW (v1.8.1 in BioEdit v.7.0.9.0. software). Indeed, before phylogenetic analysis, datasets and multiple sequence alignments were thoroughly checked to eliminate misalignments and ensure the correct framing of the coding sequences. Maximum likelihood (ML) methods were used for tree construction using full-length sequences of the S segment (1690 nucleotides). Sequence evolution was modeled using the general time reversible (GTR) + Gamma model, as determined using Model Test [35]. The best-fitting ML model according to Akaike’s information criterion was the general time-reversible + γ distribution for nucleotides, as identified by Model Test [35]. The ML trees and corresponding bootstrap support values were obtained using the online software PhyML, based on nearest neighbor exchange and subtree pruning, regrafting, branch swapping and 100 bootstrap replicates [36] (available at the ATGC bioinformatics facility: http://www.atgc-montpellier.fr/).

### Anti-RVFV antibody detection in domestic ruminants and dogs

Unfortunately, anti-RVFV antibody detection could not be carried out in wildlife because blood samples were not available. In domestic animals, RVFV-specific IgM and IgG antibodies were detected using ELISA with respectively the ID Screen® Rift Valley fever IgM Capture and ID Screen® Rift Valley fever competition multispecies kits (Innovative Diagnostics, Grabels, France) according to the manufacturer’s instructions. Diagnostic sensitivity of the IgG kit is 98% and specificity 100% [37].

Because the circulation of phleboviruses other than RVFV cannot be excluded in Gabon, a subset of randomly selected positive and negative samples was tested using the virus neutralization test (VNT), considered as the gold standard method by WOAH [38]. Briefly, duplicates of two-fold serial dilutions of sera starting from 1:5 were added to 100 TCID_50_ (50% tissue culture infectious dose) of Smithburn RVFV in 96-well microtiter plates and incubated for 1 h at 37°C. Next, 100,000 Vero cells were added to each well and the plates were incubated under 5% CO_2_ for 5–6 days at 37°C. Titers were expressed as the inverse highest dilutions giving 50% of cytopathic effect. A positive control serum was included. A serum sample with a titer of 1:10 or higher was considered seropositive.

### Statistical analysis

We analyzed small ruminant serological data from ELISA using GLM (generalized linear models), with the individual serological status as the response, and potential risk factors (species, age, gender, period of sampling) as explanatory variables. Multicollinearity among explanatory variables was assessed using variance inflation factors (VIFs). The selection of the best models was based on the Akaike information criterion (AIC). A multi-model inference approach was used for the set of models with an AIC within 2 units difference of the best model [39]. Data analyses were performed using R software version 4.3.0 [40].

## Results

### RVFV genome detection and genetic diversity

Of the 163 wildlife animals sampled along three main routes in northeastern Gabon, the RVFV-specific genome was detected in four of them (two duiker species: one sample from *Cephalophus callipygus* and three from *Philantomba monticola* (Table 2, Fig 1). After sequencing the entire S segment, phylogenetic analyses were carried out to explore their genetic relatedness with all previously published RVFV S segment nucleotide sequences. All four sequences detected in duikers clustered with a human strain of RVFV isolated in Namibia in 2004, with nucleotide identity between our sequences and the Namibian sequence ranging from 99.0 to 99.8%. This cluster is closely related to the Egyptian cluster (*i.e.* cluster A following the Grobbelaar classification [41] that also includes one strain from Zimbabwe 1978 and one strain from Madagascar 1979) (Fig 2). Viral isolation was attempted on Vero cells without success.

**Fig 2.**
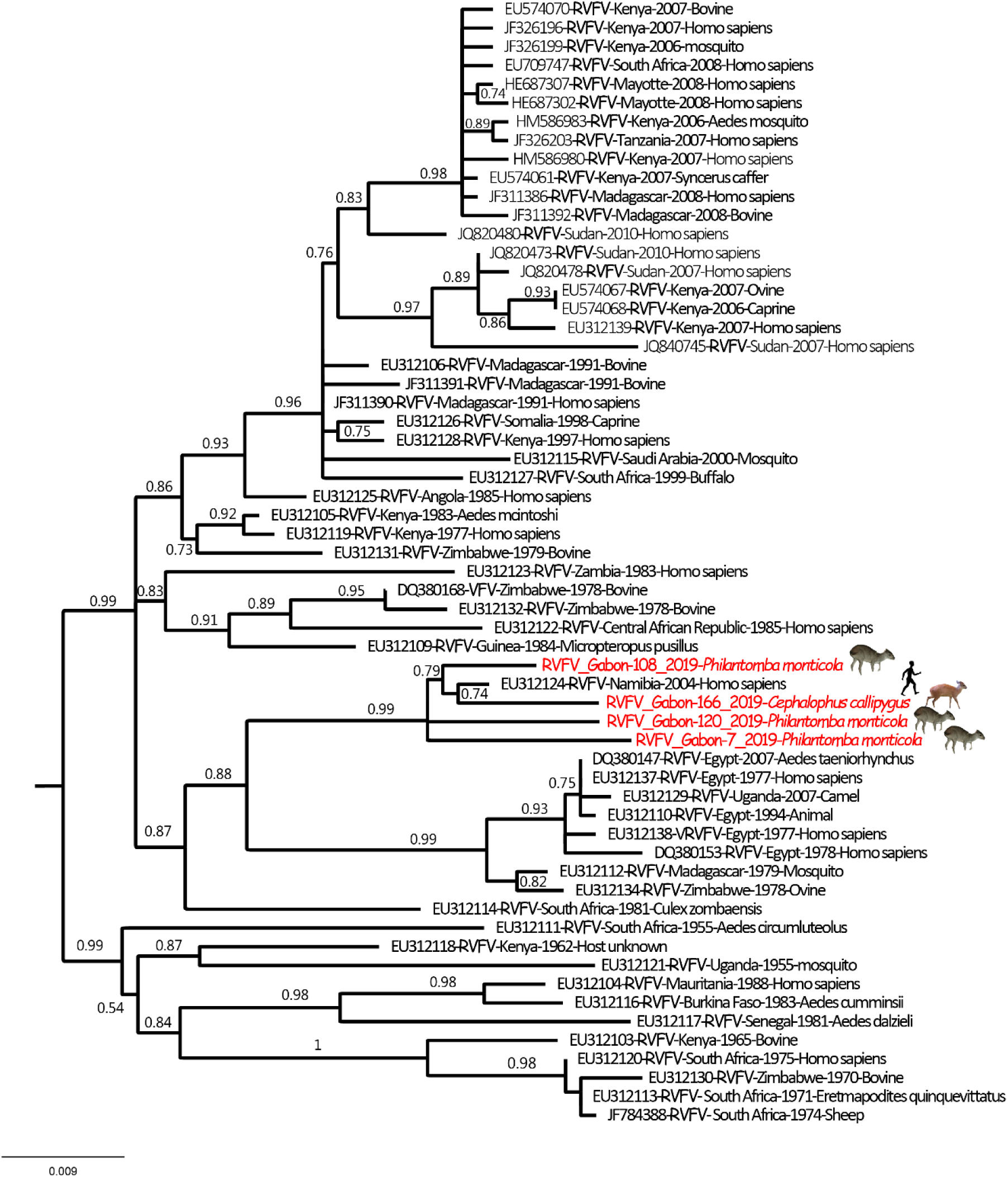
Phylogenetic tree derived from nucleotide sequence data of the entire S segment. The phylogenetic analyses were carried out after multiple alignments of the obtained sequences along with the GenBank reference sequences (including all published sequences). Maximum likelihood (ML) methods were used to construct trees based on full sequences of the S segment (1690 nt). The GenBank accession numbers for the S gene are OR528950, OR528951, OR528952, OR528953 for samples 7, 108, 120 and 166, respectively.

**Table 2.**
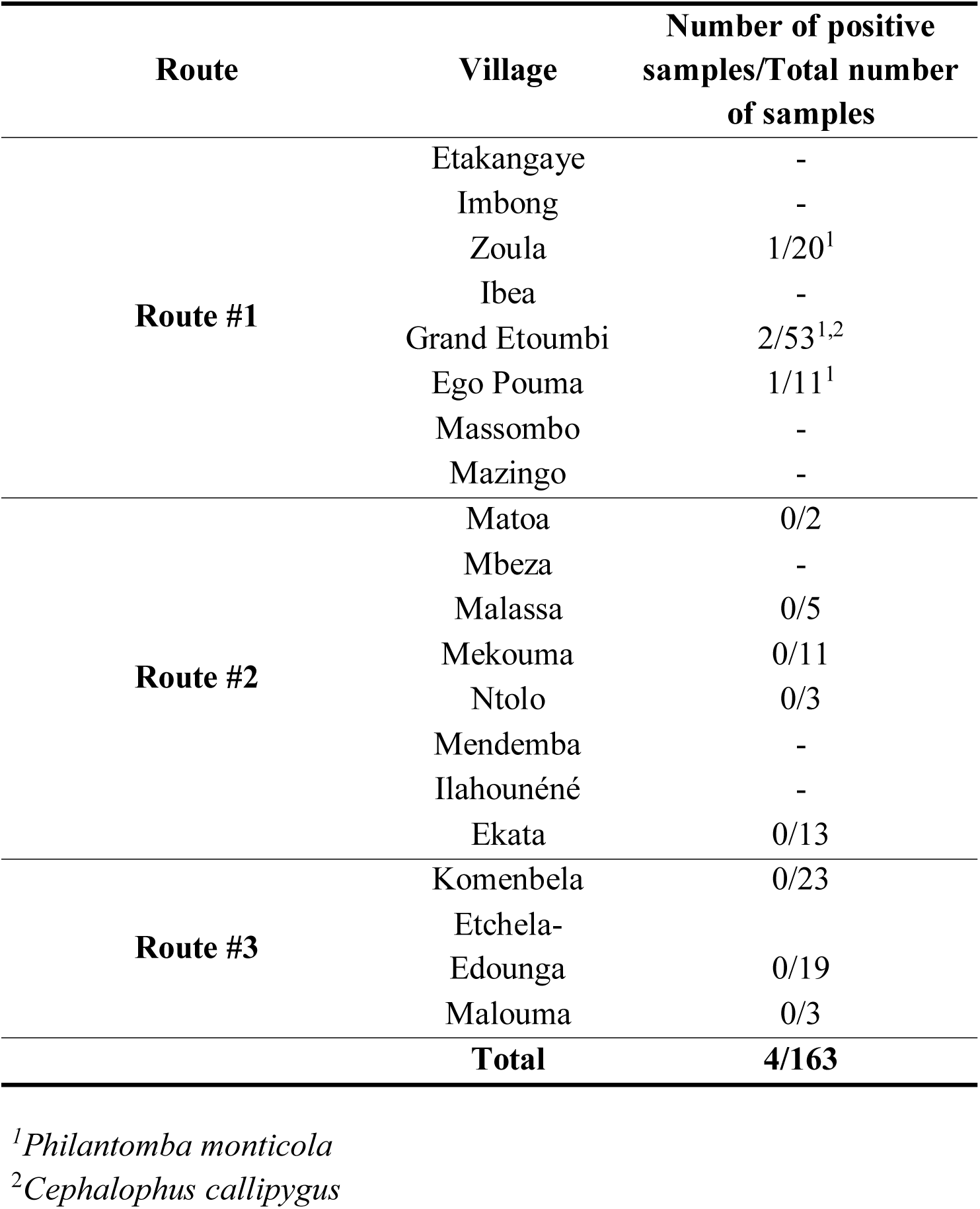
RVFV genome detection in wildlife using RT-qPCR according to sampled route and village.

### RVFV specific antibody prevalence

Following the detection of RVFV in wild duikers, a cross-sectional serological study was conducted in populations of small domestic ruminants and dogs living in villages where hunted animals were sampled. Overall, a total of 306 small ruminants (214 goats and 92 sheep) and 26 dogs (including 3 hunting dogs) were sampled and screened for RVFV specific antibodies (IgM and IgG) using ELISA. RVFV-specific IgM was not detected in any of the samples. RVFV-specific IgG antibody prevalence in livestock was 15.4% (47/306; 95% CI [11.5–19.9]) (Table 3). VNT was used to confirm samples detected highly positive by cELISA (optical density (OD) < 0.3) with 15 samples confirmed positive by VNT out of 19 tested. RVFV-specific antibodies were found in goats along both routes with similar prevalence rates (Route #1: 24.5% (23/94); Route #2: 17.5% (21/120)). Unlike goats with a seroprevalence of 20.6% (95% CI [15.4–26.6]), RVFV-specific antibodies were detected only in three sheep with a seroprevalence of 3.3% (95% CI [0.7–9.2]): Route #1: 2.6% (2/76); Route #2: 6.2% (1/16). Finally, RVFV-specific antibodies were detected in only one dog (3.8% (1/26), 95% CI [0.0– 19.6]), which was a hunting dog (Table 3).

**Table 3.**
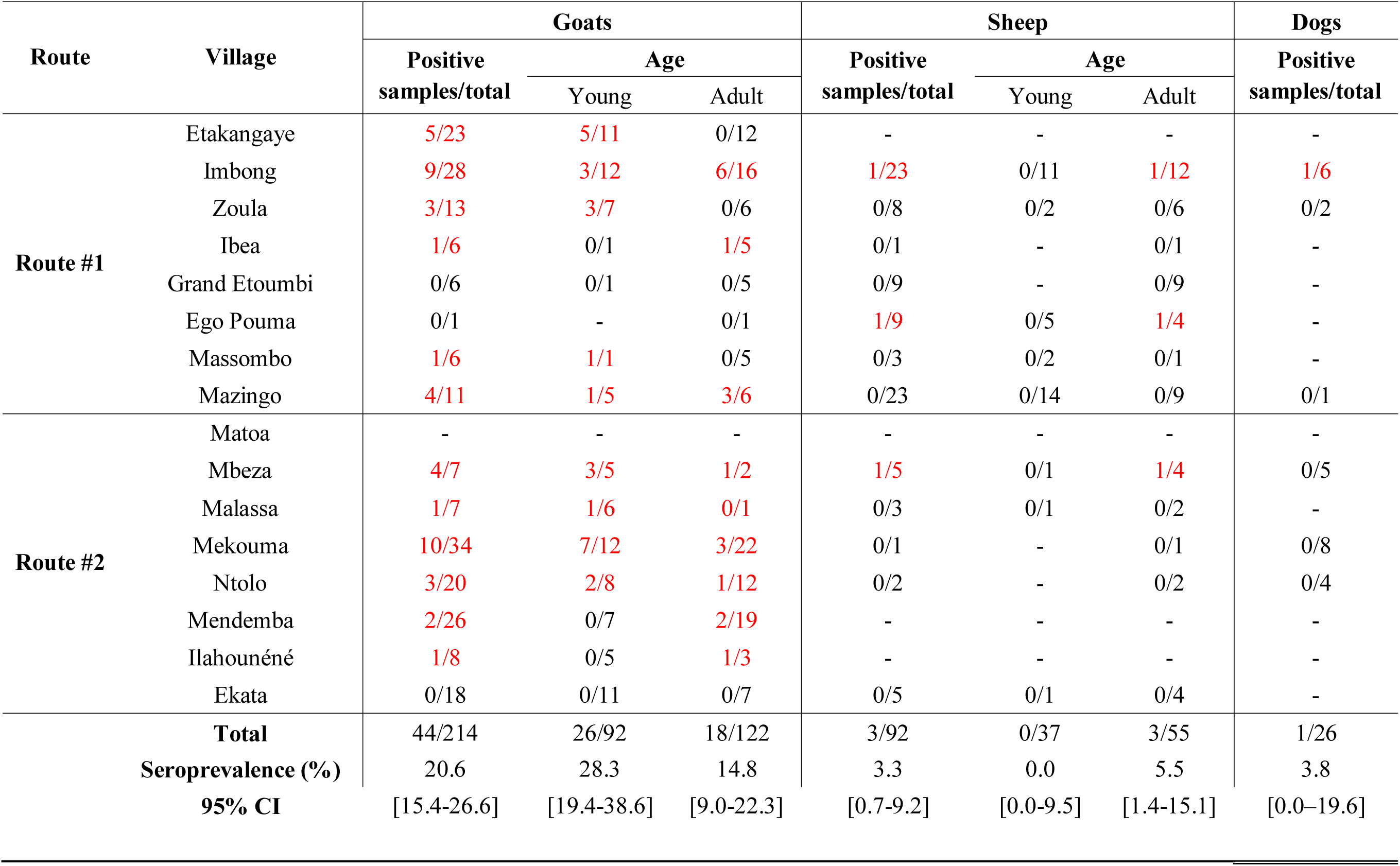
RVFV-specific IgG antibodies detection in livestock (sheep and goats) and dogs using ELISA according to sampled route and village.

Explanatory variables were not collinear (VIFs (Variance Inflation Factor) less than 2) in the full model including all explanatory variables (species, age, gender, period of sampling). According to AIC, two models were suitable for describing small ruminant seroprevalence and thus were analyzed using a multi-model inference approach. These two best models included species and period of sampling as explanatory variables. The seroprevalence in goats was significantly higher than in sheep (p<0.001; odds ratio (OR) = 8.1, 95% CI [1.4–27.1]), whereas the period of sampling was not significant using the multi-model inference.

## Discussion

Even almost 100 years after the first report of RVF, outbreaks are still difficult to anticipate and control, because the drivers of RVF endemicity are not clearly understood. The multiplicity of vertebrate hosts and mosquito species involved in the RVFV transmission cycle, the diversity of ecosystems in which RVFV occurs and the global change in human activities along with their environmental dynamics, make the entire epidemiological RVFV cycle complex and hard to determine. Although sylvatic circulation has been suspected for a long time in Central Africa, RVF rarely occurs in an epizootic form in livestock and very few clinical cases of infection have been reported in humans. The aim of this study was to investigate the circulation of RVFV in non-human vertebrate hosts living in the forest (wildlife) and villages (domestic animals) in northeastern Gabon.

### Sylvatic compartment

We here detected the RVFV genome for the first time in a forest environment in two wildlife ruminant species (duiker antelopes, *C. callipygus* and *P. monticola*) sampled in three neighboring villages in northeastern Gabon in a rainforest area. The sequencing of the S segment showed that RVFV detected in these duikers clustered with a sequence of RVFV from Namibia in 2004, closely related to the Egyptian clade A based on the Bird et al. and Grobelaar classifications [34,41]. Unfortunately, the lack of recent RVFV sequences precludes establishing links with strains circulating in Central Africa. Our results clearly demonstrate the circulation of the virus in wild animals. However, the mode of circulation of this virus remains unknown and the community of potential RVFV mosquito vectors in forest ecosystems poorly characterized, thereby requiring further investigation to model the virus transmission and maintenance. Nevertheless, several mosquito species incriminated as potential RVFV vectors (at least 14) are present in Gabon [10,42,43]. For genus *Aedes* (subgenus *Neomelaconion*), *Ae. macintoshi* has previously been reported in the country (as *Ae. lineatopennis*), as well as *Ae. palpalis* (from which the virus was previously isolated in a forest in the CAR). Interestingly, among the potential vectors present in Gabon, *Anopheles coustani* has been shown to bite wild ungulates in the forests (including *C. callipygus*), as well as six additional anopheline species (*An. carnevalei*, *An. marshallii*, *An. moucheti*, *An. obscurus*, *An. paludis*, *An. vinckei*), making these species putative candidates for the sylvatic transmission of RVFV in wild ungulates [43]. Further investigations must focus on mosquitoes that feed on wild ungulates and the Cephalophinae antelope species, they favor to determine vector candidates and are likely to shed light on sylvatic vector transmission of RVFV in Gabon.

### Domestic compartment

A cross-sectional serological study on domestic animals living in villages in the Mekambo area highlighted that goats, sheep and dogs are exposed to RVFV (overall anti-RVFV antibody prevalence of 15.4% for domestic ruminants), demonstrating its circulation in an anthropogenic environment. Most of these animals are raised locally with no history of importation or vaccination, even though a few of them come from rare imports from the Zadié Department (or from villages located on the other side of the border, in the Republic of the Congo) as gifts for weddings (dowries), deaths or religious celebrations [44]. Although our results suggest RVFV transmission at the edge of the rainforest, the origin of this circulation could also be explained by possible and rare introductions (purchases, gifts) of infected small ruminants from another area in Zadié Department or from neighboring villages in the Republic of the Congo, thus leading to virus circulation in this region.

Our study also showed that the antibody prevalence of RVFV specific antibodies was higher in goats than in sheep (20.6% *versus* 3.3%, Table 1C). To our knowledge, such a difference in seroprevalence levels observed between goats and sheep has not been reported in previous studies. However, none of them have been conducted in villages located in a forest environment, notably the recent studies carried out in the Congo Basin [18,19,23]. RVFV can be transmitted in livestock through different routes: bites of competent mosquito vectors, aerosols, contact with infected blood, body fluids and tissues of infected animals, aborted fetuses, placental membranes containing large numbers of virus particles that can either contaminate the local environment directly or infect animals [45]. In our study area, the small ruminants are not enclosed in pens and wanders around the villages. In this type of environment, goats are known to venture to the outskirts of villages readily, especially to the edge of the forest [46], according to testimonies collected from owners and villagers. Goats would be more likely exposed to forest-dwelling mosquito vectors, including those that transmit RVFV among wildlife. In contrast, sheep, which are reared around houses, are likely mainly exposed to a more domestic mosquito community. Moreover, the sheep and goats of the area may not be similarly exposed to mosquito vectors, due to qualitative and/or quantitative differences in their attractiveness to mosquitoes. Although comparative studies of goats and sheep regarding mosquito attraction are rare, some of them — undertaken in West [47] and East Africa [48,49] — suggest that there are both qualitative and quantitative differences. In Nigeria, the overall exposure of goats to mosquito bites is twice as high in goats as in sheep, but at a specific level, some mosquito species, such as *Anopheles squamosus,* incriminated as a candidate vector species during RVFV epizootics in Madagascar and Kenya [10], prefer (about 4 times as much) sheep over goats [47]. Among mosquito species involved in RVFV transmission in Kenya, *Aedes ochraceus* and *Aedes mcintoshi* seem to prefer goats over sheep, while the contrary was observed for *Mansonia uniformis* [48]. In another study from Kenya, most of the RVFV vector species (including *Ae. mcintoshi)* showed no differences in their trophic preferences between goats and sheep, although *Aedes dentatus* tended to prefer goats and *Culex pipiens* preferred sheep[49]. Nevertheless, the community of potential RVFV mosquito vectors in villages of this study area as part of the forest ecosystems remains poorly understood. It would be helpful to better document mosquito species’ blood feeding patterns in the villages of the Mekambo area, as well as to test for a possible differential host trophic preferences between goats and sheep. Excess mortality in sheep due to RVFV infection could also explain the differences in seroprevalence, but no animal owner indicated significant mortality in sheep during the sampling campaigns.

Very little data is available on RVFV circulation in dogs. To our knowledge, only one dog was reported seropositive (out of four tested using the hemagglutination-inhibition test) during the RVFV epizootic in Egypt in 1977–78 [50]. Another study reported RVFV specific antibodies using the same method in wild dogs (and none in domestic dogs) in Botswana, Kenya and South Africa [51]. However, these results could not be confirmed by VNT. Interestingly, despite our small sample size, the only seropositive dog was a hunting dog (1/3 *versus* 0/23 domestic dogs). Therefore, this dog may have been infected in the forest (via mosquito bites, or contact with an infected animal or its fluids/tissues), opening an additional opportunity for the RVFV to be introduced into the domestic compartment and subsequent transmission to domestic animals and humans: numerous anthropogenic mosquito species have opportunistic feeding habits in tropical Africa (e.g. *Culex quinquefasciatus*, *Anopheles gambiae*, *Anopheles funestus*, *Aedes albopictus*) [52–55].

### Interconnections between sylvatic and domestic compartments

The interconnections between the sylvatic and domestic compartments in a forest environment may thus be a source of zoonotic disease emergence, specifically RVFV in our case. Exploration of RVFV transmission to domestic animals in an anthropogenic environment, including the identification, the role and the blood-feeding patterns of the potential vectors, need to be explored. Comparison of viral sequences obtained from wild animals, small ruminants and dogs can help confirm whether the virus circulates between the sylvatic and domestic compartments. If there are indeed interconnections, several hypotheses need to be tested: Are there common insect vectors feeding on both wild and domestic animals? Are there overlapping areas/ecosystems where animals may be exposed to common vectors? How mobile are these animals? From which compartment does the virus emerge?

Further studies need to be carried out to understand how RVFV circulates in the forest environment of Central Africa — which is at the crossroads between West and East Africa — to investigate the sylvatic circulation of RVFV in Central African rainforests and to explore the mechanisms by which the virus shifts from its sylvatic compartment to an anthropic one, *i.e.* transmission to domestic animals and humans in villages.

This preliminary study also emphasizes the need to develop adequate event-based surveillance and control measures to limit the threat of RVF, such as awareness campaigns for villagers to report unusual deaths or abortions in domestic and wild ruminants and on the risk of RVFV infection through the manipulation of aborted fetuses, if clinical cases occur. Limiting the movement of livestock can also be proposed as a control measure. Further virological and serological dynamic surveys to investigate RVFV circulation (wet and dry seasons) in domestic animals, wildlife, hematophagous arthropods and in humans can also lead to a better understanding of RVFV circulation in the forest ecosystem of the Congo Basin.

## Acknowledgements

We thank Philippe Engandja, CIRMF, for his technical assistance. The authors thank CIRMF, IRD and OMSA for general support. We thank the people who kindly participated in our study in the different villages of the Ogooué-Ivindo province. We also thank Carolyn Engel-Gautier for scientific English language editing.

## Institutional Review Board Statement

The study was conducted in accordance with the Declaration of Helsinki and approved by the Gabonese Ministry of Health (research authorization 00093/MSP/SG/SGAQM).

## Data availability

The data are available from the corresponding author upon reasonable request.

## Author contributions

P.B., E.M.L. and G.D.M. designed the study. P.B. supervised the study conduct. P.B., G.D.M., L.B.K., T.G.M., L.H.L. I.M.M. and M.F. took part in field missions. P.B., L.B.K, T.G.M., D.G., N.N., J.V., C.P.-M., L.B., M.A.-G., N.N, MF and C.S.-S. developed the assay, performed the laboratory analyses and summarized the data in tables and figures. P.B., C.P., C.S.-S., N.D. E.M.L, and M.-M.O. analyzed the data. P.B., C.P., G.D.M. and C.C.-S. wrote the manuscript, and all authors contributed to the text and approved the final version of the manuscript.

